# Response to comment on ‘Initiation of chromosome replication controls both division and replication cycles in *E. coli* through a double-adder mechanism’

**DOI:** 10.1101/2020.08.04.227694

**Authors:** Guillaume Witz, Thomas Julou, Erik van Nimwegen

## Abstract

Last year we published an article (***Witz et al***., 2019) in which we used time-lapse microscopy in combination with microfluidics to measure growth, division and replication in single *E. coli* cells on the one hand, and developed a new statistical analysis method to calculate the ability of different cell cycle models to capture the correlation structure observed in the data on the other hand. This led us to propose a new model of cell cycle control in *E. coli* which we called the double-adder model.

Recently Le Treut et al. published a comment (***Le Treut et al***., 2020) on our article which made a number of highly critical claims, including allegations that our own data support a different model than the one we proposed, and that our model cannot reproduce the ‘adder phenotype’ observed in the data. We here show that all these allegations are false and based on basic analysis errors. Although our focus is on explaining the errors in the analysis of Le Treut et al, we have attempted to make the presentation of interest to a broader scientific audience by discussing the issues in the context of what our current understanding is of the bacterial cell cycle, and to what extent recent data either support or reject various proposed models.

## Introduction

Le Treut et al. recently published a comment on our article ***Witz et al***. (***2019***) and submitted the same comment for publication to eLife (***Le Treut et al***., 2020). The journal invited us to write a formal response and based on review of the editors and an external expert of both the comment of Le Treut et al. and our response, the journal decided to reject the comment of Le Treut et al. However, they strongly suggested we publish our response on bioRxiv to make sure it is available to the public.

The comment of Le Treut et al. severely criticized our work. In particular, Le Treut et al. claim that our data support a different model than the one we propose, that the correlation analysis that we introduced in any case cannot distinguish different models, and that our model is at odds with the ‘adder phenotype’ observed in the data. All these claims are false. As we will show below, the correlation analysis results that Le Treut et al. report are all meaningless because of basic errors in the application of this method. In addition, the claim that our model fails to reproduce the adder phenotype is based on comparing the experimental data not with our actual model, but with a mutilated version that the authors concocted by altering the simulation code that we provided.

Since all the results presented by Le Treut et al. are based on faulty analysis, we cannot see anything constructive in their contribution, and we fear that the main result of their comment will be to muddy the waters regarding our current understanding of the bacterial cell cycle. While data from the additional experiments that Le Treut et al. report might well provide a valuable addition to the literature, their analysis of this data is equally faulty, so that it is currently unclear what additional insights might be gained from this data.

## Methods and Results

### Division-centric? Replication-centric? What exactly are we talking about?

The main impetus behind the comment of Le Treut et al. seems to have been a perceived contradiction between our results and the results they presented in ***Si et al***. (***2019***), and much of their discussion concerns comparison of ‘their’ division-centric and ‘our’ replication-centric model of cell size homeostasis. However, in the absence of specifying concrete models, these terms are ambiguous and we suspect that many readers may easily lose track of what the actual differences are between these models, as well as to what extent the data presented in ***Si et al***. (***2019***) and ***Witz*** et al. (***2019***) are even at odds or supporting one model over the other. We will thus start by giving a short overview of what precisely these models are and to what extent different data either support or reject them. Notably, given the large variety of cell cycle models that have been proposed in the literature, and the emphasis that Le Treut et al. put on contrasting their model with ours, the reader may be surprised to learn that these models are actually almost identical, differing in only one aspect.

To start from a historical reference point, the influential Helmstetter-Cooper model assumes that replication is initiated at a fixed critical cell size and that a fixed time elapses between replication initiation and division (***Cooper and Helmstetter***, 1968). Using a combination of microfluidics with time-lapse microscopy, in recent years researchers have been exploring to what extent the observed correlation structure of the fluctuations in the sizes and times at birth, initiation and division in single cells are consistent with such models of the cell cycle. In particular, if a model assumes that the cell cycle control mechanism acts through constraining a particular variable X, then one would expect single-cell fluctuations in X to be independent of fluctuations in other quantities.

For example if, as the Helmstetter-Cooper model assumes, initiation is triggered when the cell reaches a critical size, one would expect fluctuations in the size at initiation to be independent of fluctuations in the size at birth. However, recent works, including ours (Fig. 2A), show that there is a clear positive correlation between size at birth and initiation (***Micali et al***., 2018a; ***Witz et al***., 2019). In this way, a particular model of the cell cycle can be falsified by showing that the observed correlations are at odds with predictions of the model.

An alternative model of the replication cycle assumes replication initiation is controlled by an inter-initiation adder, i.e. that cells add a fixed volume per origin between consecutive replication events. Indeed, both in ***Si et al***. (***2019***) and in our study (***Witz et al***., 2019) it was observed that the added volume per origin fluctuates independently from the size at initiation, supporting this model. However, it is important to note that this observed independence does not in itself prove that replication must be controlled by an inter-replication adder mechanism, i.e. other models may well also predict that there is no correlation between size at initiation and added volume between initiations.

As a case in point, the so called ‘adder phenotype’ corresponds to the observation that the volume added between birth and division is approximately uncorrelated with the cell’s size at birth (***Amir***, 2014; ***Osella et al***., 2014; ***Campos et al***., 2014; ***Taheri-Araghi et al***., 2015). Although this observation can of course be explained by an inter-division adder model, i.e. assuming that cells directly constrain the volume added between birth and division, other models also predict the adder phenotype. For example, even before the papers demonstrating the adder phenotype appeared, theoretical work by Amir showed that such an adder phenotype is also predicted by a model that combines an inter-initiation adder with a fixed time between replication and division, i.e. as in the Helmstetter-Cooper model (***Amir***, 2014; ***Ho and Amir***, 2015; ***Amir***, 2017). In our work we thus stressed that, in order to meaningfully compare the evidence that the single-cell data provide for one or another model, one should compare the *full* correlation structure of the data with the predictions of the different models.

We show in ***Witz et al***. (***2019***) that, while the data are consistent with an inter-initiation adder model, the time between replication initiation and division in fact correlates with growth-rate (Fig. 3A), thereby falsifying a model that assumes division is controlled by a mechanism that acts on the time between initiation and division. Instead, our analysis shows that in the model which is most consistent with the full correlation structure of our data, division is controlled by an adder mechanism that runs from replication-initiation to division. In this model, each time replication is initiated an adder is started for each replicated origin, and each of these adders trigger a subsequent division event when a total critical amount of volume has been added. Importantly, we show that this model also reproduces the ‘adder phenotype’, i.e. that added volume between birth and division is approximately uncorrelated with the cell’s size at birth (Fig. 5A).

Thus, the model that we propose assumes that an inter-initiation adder controls replication initiation, that division is controlled by an adder running from replication-initation to division, and that growth-rate fluctuates independently from the replication and division control. The model proposed in the comment of Le Treut et al. is almost identical. First, the model of Le Treut et al. also assumes that replication initiation is controlled by an inter-initiation adder and in this sense replication control is thus ‘replication-centric’ in both models. Second, both models assume that growth-rate is an independent variable whose fluctuations are uncoupled from the control of the sizes at which replication and division occur. Third, both models also agree that division is controlled by an adder mechanism. The only difference is that, while our model assumes this adder is initiated at replication initiation, the model that Le Treut et al. propose assumes the adder is initiated at birth. For future reference we will refer to these models as the Replication Double Adder (RDA) and Independent Double Adder (IDA) models.

### What does the evidence say?

Although Le Treut et al claim that the IDA model has been ‘revealed’ in ***Si et al***. (***2019***), a reader might be excused for not coming away with this impression from reading this paper. ***Si et al***. (***2019***) show that cells exhibit ‘adder phenotypes’ for both the replication and division cycles, i.e. that the inter-initiation added volume is approximately uncorrelated with the size at initiation, and that the added volume between birth and division is approximately uncorrelated with the size at birth. Then, by applying various perturbations, including massive time-dependent perturbations in the levels of key cell cycle proteins, ***Si et al***. (***2019***) show that the correlations that define the replication adder and division adder phenotypes can be separately perturbed. However, instead of putting forward the IDA model, in ***Si et al***. (***2019***) the authors develop a complex variant of the Helmstetter-Cooper model which allows for an arbitrary correlation structure between all variables in the model, as well as correlation across generations, without discussing how these complex correlations could be implemented mechanistically.

In fact, we see no reason why any of the results in ***Si et al***. (***2019***) would favor the IDA over the RDA model. Given that the replication and division cycles are ultimately implemented by mechanisms that involve separate molecular players, one would generally expect that it is possible to separately perturb the two adder phenotypes. In order to predict how either the IDA or RDA models would respond to the perturbations applied in ***Si et al***. (***2019***) one would need much more concrete models of how these adder mechanisms are implemented biophysically. In any case, no attempt is made in ***Si et al***. (***2019***) to evaluate to what extent their data favor one model over the other.

In contrast, we did explicitly investigate whether the IDA model could also explain our data. Simulations of the IDA model showed that, while this model reproduces most of the statistics observed in our data, it clearly fails in one critical respect (Appendix 2 Figure 2 of ***Witz et al***. (***2019***)). That is, the IDA model predicts a strong negative correlation between the size at replication-initiation and the size added between initiation and division. However, no such correlation is observed in the data. Even though we thus specifically identified a correlation for which the IDA and RDA models make different predictions, and showed that our data clearly favor the RDA over the IDA model, Le Treut completely ignore this observation, never comment on it, and make no attempt to investigate it on their own data. This is especially remarkable given that Le Treut et al. go as far as to claim, in direct contradiction to these results, that our own data in fact favor the IDA model over the RDA model. We next discuss why those claims are based on faulty analysis.

### The correlation analysis and independence measure *I* apply only when the events of each cell cycle are independent of other cell cycles

In ***Witz et al***. (***2019***) we developed a correlation analysis method to systematically compare a large class of cell cycle models on our largest, automatically curated, dataset, which was for cells growing in minimal media with glycerol. To rigorously compare how well different cell cycle models can explain a dataset, one would generally have to formulate concrete likelihood functions that calculate the probability of all the observed data given each model, i.e. all times and sizes at birth, replication initiation, and division, for all observed lineages of cells. Depending on the growth conditions and models one wants to consider, such likelihood functions will typically contain non-trivial dependencies between events across multiple cell cycles.

However, we took advantage of the fact that, in the growth conditions we studied, cell cycles do not overlap, and considered a class of models for which, given the state of the cell at the start of its cell cycle, all events in each cell cycle are independent of the events in other cell cycles. In particular, the class of models we considered are defined by the starting point of the cell cycle, which may be at replication-initation or birth, and by a set of variables that are being independently constrained. For example, for the Helmstetter-Cooper model each cell cycle starts at replication-initiation and is defined by the growth-rate, the size at initiation, and the time between initiation and division. Similarly, in the RDA model the cell cycle also starts at initiation and is defined by growth-rate, the added volume between initiations, and the added volume between initiation and division. Given a model, the data of each cell cycle are defined by the state of the cell at the start of the cell cycle, and the values of the variables that the model assumes are independently constrained. As discussed in ***Witz et al***. (***2019***), in this situation the fit of each model to the data can be quantified by the determinant *I* of the correlation matrix. We find that, of all models considered, the RDA model has the highest value of *I* and we subsequently showed, using simulations, that the RDA model also fits the data from experiments at higher growth rates.

However, as explained above, this correlation analysis crucially relies on the fact that cell cycles are not overlapping. As soon as cells grow so fast that cell cycles overlap, events within one cell cycle depend on events that occurred one or more cell cycles in the past, and a more complex correlation analysis would be required. In addition, for the IDA model there are two adders running concurrently, so that there is there is no single event at which the cell cycle can be considered to start. Consequently, the correlation analysis presented in our paper cannot be applied to the IDA model *even at slow growth*, because events within one cell cycle depend on at least one event from a previous cell cycle. Indeed, exactly for this reason we did not attempt to calculate an *I* value for the IDA model, but instead highlighted one key correlation for which the RDA and IDA models make different predictions, as we mentioned above.

In hindsight we realize that we should have explained more clearly that the *I* values of our correlation analysis can only be meaningfully calculated when all cell cycles are non-overlapping and for models for which the events in each cell cycle are independent of the events in other cell cycles. Le Treut et al. have clearly not realized this limitation and most of the results in their comment consist of comparing *I* values for the RDA and IDA models, including on datasets with overlapping cell cycles. However, since one cannot meaningfully calculate an *I* value for the IDA model, nor for growth conditions with overlapping cell cycles, all these reported results are essentially meaningless.

### The correlation analysis does require 4 and not 3 variables

However, apart from these problems, the *I* values that Le Treut et al. report are also meaningless because of an additional error. Instead of following our method and calculating the determinant of a 4 by 4 correlation matrix of 4 variables, Le Treut et al. claim that it suffices to calculate the determinant of only a 3 by 3 correlation matrix. In particular, they argue this suffices because cell cycle models can be defined using 3 variables only. While it is true that 3 variables may be enough to define a cell cycle model, it is not enough to fully characterize the correlation structure of the cell cycles.

To illustrate this we will use a simplified example by ignoring the replication-cycle, and considering only the division cycle running from birth to division. The measured variables for a cell cycle will then consist of the length at birth *L*_*b*_, the length at division *L*_*d*_ and the time *T* between birth and division. In our correlation analysis, each possible model describes the events of each cell cycle by the state of the cell at birth and two additional variables that control the timing and size at division, and then calculates the determinant *I* of the 3 by 3 correlation matrix of these variables. For example, the top row of Fig. 1 shows the correlation matrices and independence values *I* for the adder model and for a sizer model as calculated from our experimental data in slow growth conditions (M9-glycerol) used in Fig.6 of (***Witz et al***., 2019). That is, for the adder model each cell cycle is characterized by size at birth *L*_*b*_, added length *dL* = *L*_*d*_ − *L*_*b*_, and the growth-rate *λ*. Similarly, for a sizer model each cell cycle is characterized by size at birth *L*_*b*_, size at division *L*_*d*_, and the growth-rate *λ*. As expected, the *I* values indicate that the adder model is by far preferred over the sizer model.

**Figure 1.**
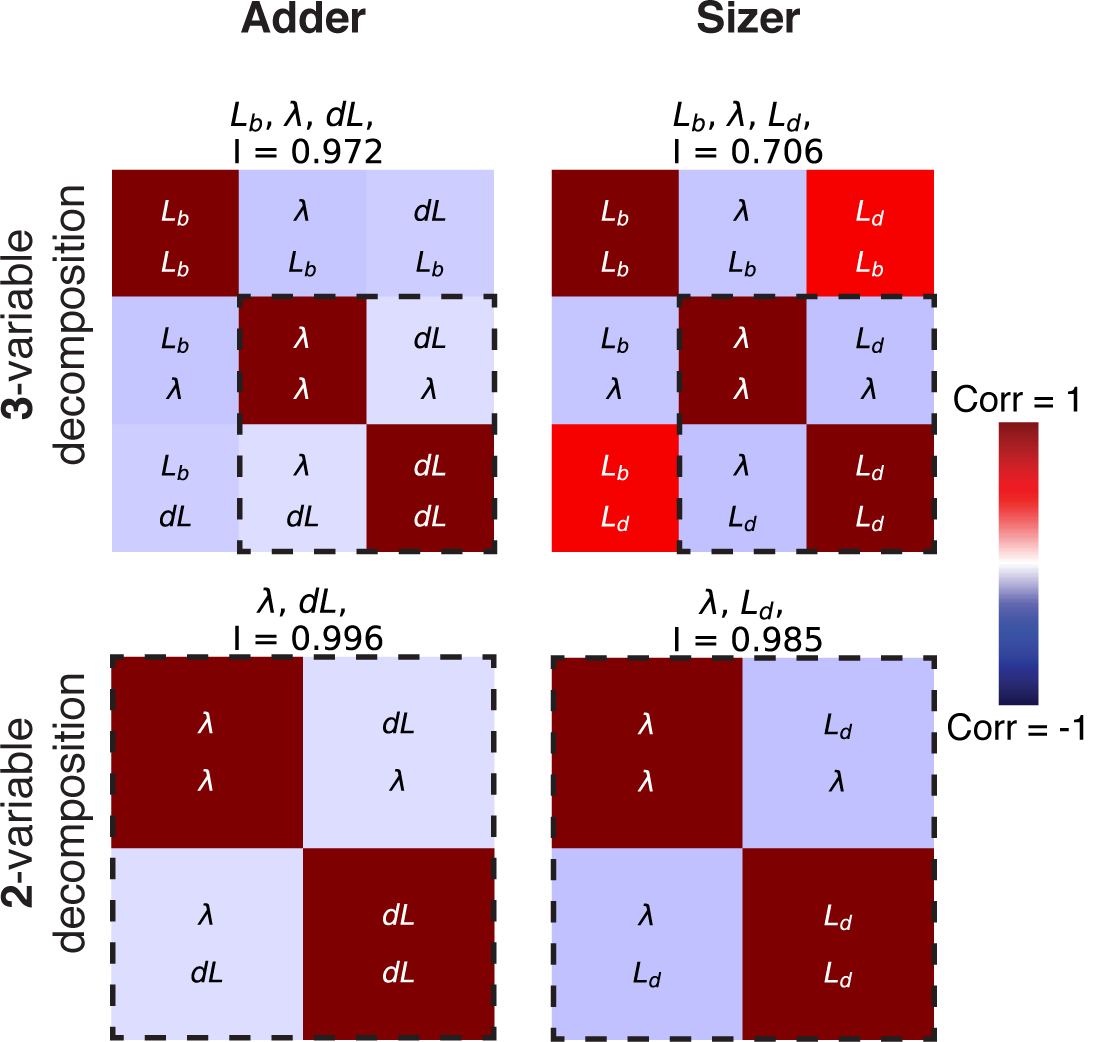
Correlation analysis for the division cycle. The top two panels show the correlation matrices for the decomposition corresponding to the adder model which uses cell length at birth *L*_*b*_, growth-rate *J* and added length *dL* (left) and the decomposition for a sizer model which uses length at birth *L*_*b*_, growth-rate *J*, and length at division *L*_*d*_ (right). Note that the sizer has a much lower *I* value because of the correlation between size at birth and division. However, if we drop the size at birth *L*_*b*_ from the correlation matrix, as Le Treut at all suggest we can, we obtain the two correlation matrices in the bottom panels which both have virtually identical independence value *I*, since the crucial correlation between size at birth and division is now missing.

However, one might argue that only 2 variables are needed to define the adder and sizer models, i.e. for each cell cycle we can draw the growth-rate *λ* and either *dL* or *L*_*d*_ from an independent distribution. However, as shown in the panels of the bottom row of Fig. 1, if we leave the initial state *L*_*b*_ out of the correlation matrix, and calculate *I* values for the 2 by 2 correlation matrices, there is no longer any meaningful difference between the *I* values of the two models. That is, the crucial evidence for the adder model is in the correlation between the sizes at birth and division and this is lost when the initial size *L*_*b*_ is left out of the correlation matrix. While it is true that one can think of the adder model as only specifying *λ* and *dL* of each cell cycle, once we leave out *L*_*b*_ the size at birth of one cell depends on the values of *dL* in all its ancestor cells, thereby coupling the data from different cell cycles together. Since the correlation analysis requires treating each cell cycle as independent, it is thus crucial to include the size at birth.

Unfortunately, Le Treut et al. have also not recognized this problem, and consistently calculate *I* values for 3 by 3 correlation matrices. These *I* values are thus meaningless for at least two separate reasons.

### Since Le Treut et al’s correlation analysis is invalid, discussions about the role of slow growth, the precise strain, or the origin tracking method are all moot

Le Treut et al. spend a considerable portion of their comment comparing *I* values of the RDA and IDA models for datasets from different experiments, showing that the *I* values they calculate slightly prefer either the RDA or IDA model depending on whether one considers slow growth conditions, what precise *E. coli* lab strain is used, whether one supplements uracil in the growth media, and whether one uses FROS or *dnaN-YPet* to track replication initiation. Since, as we just explained, these *I* values result from a faulty analysis, the small differences that are observed are meaningless and thus not worthwhile discussing.

However, even *if* we were to assume that the *I* values were meaningful, the logic of Le Treut et al’s discussion of these results seems rather confused. That is, Le Treut et al. suggest that the fact that RDA model is preferred over the IDA model on some of their own datasets is an artefact resulting either because of slow growth, or because of a growth defect due to a *rph-1* mutation that requires uracil supplementation to overcome, or due to using FROS. However, as we already mentioned above, Le Treut et al. also claim that our own data prefer the IDA over the RDA model, and those data were obtained using a strain with the same *rph-1* mutation as strain MG1665, without supplementing uracil, and using FROS instead of *dnaN-YPet*. This thus directly contradicts that these details would cause the RDA model to be ‘artificially’ preferred.

### Independence values *I* can only be meaningfully calculated for valid cell cycle decompositions

Even though most of the results that Le Treut et al. present consist of comparison of *I* values, in the last part of their comment they go on to claim that these *I* values are in fact too insensitive to meaningfully distinguish between different cell cycle models. To support this claim, they start from their set of 18 possible variables that can be used in cell cycle decompositions and calculate *I* values for *all* 816 possible subsets of 3 variables, showing that these *I* values have a smooth distribution, and noting also that the IDA and RDA models rank only 57th and 88th, respectively. While we have some understanding for the previous mistakes made by Le Treut et al., i.e. mistakenly assuming that it was meaningful to calculate *I* values for the IDA model and in situations with overlapping cell cycles, and mistakenly assuming that a 3 by 3 correlation matrix would suffice, we are frankly astonished that they do not seem to realize that the vast majority of these 816 subsets cannot specify cell cycle models at all. For example, size at birth *L*_*b*_, size at division *L*_*d*_, and added size *dL* = *L*_*d*_ − *L*_*b*_ would be one of the 816 possible subsets. However, this subset does not specify anything about either replication initiation nor anything about the timing of the cell cycle events. It is of course completely meaningless to calculate *I* values for random subsets of variables like this. Moreover, we explicitly showed in our paper (Fig. 7A), that if one calculates *I* values only for meaningful decompositions (listed in Fig. 7S3), the top RDA decomposition clearly stands out, and all division-centric decompositions perform poorly.

### The RDA model does reproduce the observed ‘adder phenotype’

Le Treut et al. also claim that the RDA model cannot reproduce the adder phenotype, i.e. that the added volume between birth and division is approximately independent of the volume at birth. To support this they analytically calculate the expected correlation coefficient between size at birth and division for an idealized version of the RDA model and show that this predicted correlation, while positive, is less than the correlation of 1/2 that would be predicted for a simple adder model. However, in our paper we explicitly showed through simulations of the RDA model that it *does* reproduce the ‘adder phenotype, i.e. see Fig. 5A of ***Witz et al***. (***2019***).

So how is this possible? The reader will have to turn to the materials and methods of the comment of Le Treut et al. to discover that, although they used our simulation code to simulate the RDA model, they in fact modified the code so as to remove small stochastic fluctuations in the sizes of the two daughters at division. That is, we observed in the experimental data that a mother cell of size 2*L* does not produce two daughters of perfectly equal size *L*, but daughters of sizes *L* + *δ* and *L* − *δ* where *E*/*L* has a standard-deviation of about 0.06, i.e. six percent fluctuations in the sizes of the two daughters. We incorporated this asymmetry into our simulations because it significantly improved the fit of the observed and predicted variation in cell size at birth, but it also improves the ‘adder phenotype’, i.e. the fit between the predicted and observed correlations in added volume *dL* and size at birth *L*_*b*_ of Fig. 5A. Indeed, in the Materials and Methods of their comment Le Treut et al. in fact admit that a mismatch between model and data is only observed when this asymmetry at division is removed.

We cannot understand why Le Treut et al. consider the fact that a simplified version of the RDA model (i.e. without asymmetry at division) predicts a correlation coefficient of less than 1/2 is considered an inherent flaw of the model. Whether the RDA model makes the exact same predictions as a perfect adder model is irrelevant. The only thing that matters is whether the model fits the data. Indeed, while the perfect adder predicts precisely zero dependence between the size at birth *L*_*b*_ and the added size *dL*, the RDA model predicts a weak negative correlation. In fact, a careful look at Fig. 5A of ***Witz et al***. (***2019***) shows that the experimental data appear to exhibit a small negative correlation between *L*_*b*_ and *dL* as well. Moreover, such weak negative correlations can also be seen for the data shown in Fig. 3B and 4B of ***Si et al***. (***2019***).

As noted earlier, several studies have shown that there is a clear positive correlation between size at birth *L*_*b*_ and size at initiation *L*_*i*_ (e.g. Fig 2A and 5B of ***Witz et al***. (***2019***)), rejecting models that assume a critical initiation size, because such models predict no correlation between *L*_*b*_ and *L*_*i*_. It is thus noteworthy that a simple analytical calculation shows that, without asymmetry at division, the IDA would *also* predict no correlation between the size at birth *L*_*b*_ and size at initiation *L*_*i*_. However, when we simulated the IDA model including asymmetry at division, a weak positive correlation between *L*_*b*_ and *L*_*i*_ appeared (Fig. 2 of Appendix 2 ***Witz et al***. (***2019***)). Although this correlation is clearly weaker than what is observed in the data, we considered the difference too small to reject the IDA model on that basis, and instead focused only on the clear mismatch in correlation between size at initiation and added size between initiation and division.

Finally, we think it is instructive to compare the amount of concern that Le Treut et al. express regarding the small difference in the precise correlations predicted by the RDA and perfect adder models, with the concern for accuracy in such matters that these authors exhibit in ***Si et al***. (***2019***). For example, whereas in ***Witz et al***. (***2019***) we make sure to display experimentally observed correlations using full scatters, in ***Si et al***. (***2019***) correlations are only shown using binned data, and these binned data are plotted using symbols that are so large that data from different experiments completely obscure each other, making it virtually impossible to see what precise correlations the data actually exhibit, e.g. see Figs. 1D and 7B of ***Si et al***. (***2019***). Moreover, in ***Si et al***. (***2019***) the authors routinely describe observed correlations as ‘sizers’ or ‘adders’ when the observed correlations clearly deviate substantially from what sizer and adder models would predict. For example, Fig 3B of ***Si et al***. (***2019***) purports to show that, under the perturbations in question, the relationship between size at initiation and inter-initiation added size behaves as a ‘sizer’, while the division cycles still show adder behavior, i.e. that the added length between birth and division is independent of birth size. However, the negative correlation between inter-initiation added size and initiation size is clearly less steep then a sizer would predict, and there is also clearly a weak negative trend between size at birth and added size (as an RDA model would in fact predict). Very similar comments apply to Fig. 4B of ***Si et al***. (***2019***) where the authors call a variety of clearly different slopes all ‘sizer’ whereas a clearly still negative but weaker slope is called an ‘adder’. The very weak negative correlation in Fig. 7B, which the authors call a ‘sizer, is also far from what a sizer would actually predict.

## Conclusions

It has not been a pleasant exercise to write this response. We more than welcome post-publication review and feedback on our work, but we struggle to find anything constructive in the comment of Le Treut et al, and it is hard for us to understand what motivated these authors to attack our work so vehemently and unreasonably. This is especially surprising to us given that the IDA model that they champion in their comment is so close to the RDA model that we proposed in our paper. In fact, in ***Si et al***. (***2019***) these authors repeatedly state that the main result of their experiments and analyses is that the division adder phenotype is a direct consequence of: (1) (threshold) accumulation of division initiators and precursors to a fixed threshold number per cell and (2) (balanced biosynthesis) their production is proportional to the growth of cell volume under steady-state condition. However, both of these points apply equally well to the initiation-to-division adder of the RDA model proposed in ***Witz et al***. (***2019***). As such, we do not understand why Le Treut et al. even feel that the results presented in ***Witz et al***. (***2019***) are at odds with theirs. That is, we do not see any result in ***Si et al***. (***2019***) that suggests that the division adder must be initiated at birth rather than at replication-initiation.

Finally, we would like to stress that we do not believe that these relatively simple models give anywhere near a full picture of bacterial cell cycle control. On the contrary, it seems likely to us that the evolutionary process has ‘designed’ cell cycle controls that involve multiple complementary and redundant mechanisms including multiple check points. Moreover, different mechanisms may be more or less dominating in different species or within different conditions. For example, although both the IDA and RDA models do not specify any direct coupling between the replication and division adders, we of course know that couplings must exist. For example, we know that under particular stress conditions, such as when cells filament as a result of DNA damage, additional checkpoint mechanisms clearly come into play. In our simulations of the IDA model we observed a small fraction of cell cycles where the division adder was triggered before replication had even been initiated (let alone finished), and cells undoubtedly have control mechanisms to avoid this. Indeed, ***Micali et al***. (***2018b***) recently introduced a family of models that combine concurrently running processes that separately control the replication and division cycles with an explicit check point mechanism and, as we pointed out in ***Witz et al***. (***2019***), it is plausible that such models could in principle also fit our data, provided that their parameters are precisely tuned. In summary, we feel that models such as our RDA model should only be considered as useful starting points for further exploration.

## Notes

### Competing Interest Statement

The authors have declared no competing interest.

## References

Amir A. Cell Size regulation in bacteria. Physical Review Letters. 2014 May; 112(20):208102. doi: https://doi.org/10.1103/PhysRevLett.112.208102.

Amir A. Is cell size a spandrel? eLife. 2017 01; 6. doi: https://doi.org/10.7554/eLife.22186.

Campos M, Surovtsev IV, Kato S, Paintdakhi A, Beltran B, Ebmeier SE, Jacobs-Wagner C. A constant size extension drives bacterial cell size homeostasis. Cell. 2014; 159(6):1433–1446. doi: https://doi.org/10.1016/j.cell.2014.11.022.

Cooper S, Helmstetter CE. Chromosome replication and the division cycle of Escherichia coli B/r. Journal of Molecular Biology. 1968 Feb; 31(3):519–540. doi: https://doi.org/10.1016/0022-2836(68)90425-7.

Ho PY, Amir A. Simultaneous regulation of cell size and chromosome replication in bacteria. Frontiers in Microbiology. 2015; 6:662. doi: https://doi.org/10.3389/fmicb.2015.00662.

Le Treut G, Si F, Li D, Jun S. Comment on ‘Initiation of Chromosome Replication Controls Both Division and Replication Cycles in E. Coli through a Double-Adder Mechanism’. bioRxiv. 2020; 2020.05.08.084376. doi: https://doi.org/10.1101/2020.05.08.084376.

Micali G, Grilli J, Marchi J, Osella M, Cosentino Lagomarsino M. Dissecting the control mechanisms for DNA replication and cell division in *E. coli*. Cell Reports. 2018; 25(3):761–771.e4. doi: https://doi.org/10.1016/j.celrep.2018.09.061.

Micali G, Grilli J, Osella M, Cosentino Lagomarsino M. Concurrent processes set *E. coli* cell division. Science Advances. 2018; 4(11):eaau3324. doi: https://doi.org/10.1126/sciadv.aau3324.

Osella M, Nugent E, Cosentino Lagomarsino M. Concerted control of *Escherichia coli* cell division. PNAS. 2014; 111(9):3431–3435. doi: https://doi.org/10.1073/pnas.1313715111.

Si F, Le Treut G, Sauls JT, Vadia S, Levin PA, Jun S. Mechanistic origin of cell-size control and homeostasis in bacteria. Current Biology. 2019 06; 29(11):1760–1770. doi: https://doi.org/10.1016/j.cub.2019.04.062.

Taheri-Araghi S, Bradde S, Sauls JT, Hill NS, Levin PA, Paulsson J, Vergassola M, Jun S. Cell-size control and homeostasis in bacteria. Current Biology. 2015; 25(3):385–391. doi: https://doi.org/10.1016/j.cub.2014.12.009.

Witz G, van Nimwegen E, Julou T. Initiation of chromosome replication controls both division and replication cycles in E. coli through a double-adder mechanism. eLife. 2019 11; 8. doi: https://doi.org/10.7554/elife.48063.

